# On the importance of skewed offspring distributions and background selection in viral population genetics

**DOI:** 10.1101/048975

**Authors:** Kristen K. Irwin, Stefan Laurent, Sebastian Matuszewski, Séverine Vuilleumier, Louise Ormond, Hyunjin Shim, Claudia Bank, Jeffrey D. Jensen

## Abstract

Many features of virus populations make them excellent candidates for population genetic study, including a very high rate of mutation, high levels of nucleotide diversity, exceptionally large census population sizes, and frequent positive selection. However, these attributes also mean that special care must be taken in population genetic inference. For example, highly skewed offspring distributions, frequent and severe population bottleneck events associated with infection and compartmentalization, and strong purifying selection all affect the distribution of genetic variation but are often not taken in to account. Here, we draw particular attention to multiple-merger coalescent events and background selection, discuss potential mis-inference associated with these processes, and highlight potential avenues for better incorporating them in to future population genetic analyses.

## Introduction

Viruses appear to be excellent candidates for studying evolution in real time; they have short generation times, high levels of diversity often driven by very large mutation rates and population sizes (both census and effective), and they experience frequent positive selection in response to host immunity or antiviral treatment. However, despite these desired attributes, standard population genetic models must be used with caution when making evolutionary inference.

Firstly, population genetic inference is usually based on a coalescence model of the Kingman type, under the assumption of Poisson-shaped offspring distributions where the variance equals the mean and is always small relative to the population size; consequently, only two lineages may coalesce at a time. In contrast, viruses have highly variable reproductive rates, taken as rates of replication; these may vary based on cell or tissue type, level of cellular differentiation, or stage in the lytic/lysogenic cycle (Knipe and Howley, 2007), resulting in highly skewed offspring distributions. This model violation is further intensified by the strong bottlenecks associated with infection and by strong positive selection (Neher and Hallatschek, 2013). Therefore, virus genealogies may be best characterized by *multiple merger* coalescent (MMC) models (e.g, Pitman, 1999; Sagitov, 1999; Donnelly and Kurtz, 1999; Schweinsberg, 2000; Möhle and Sagitov, 2001; Eldon and Wakeley, 2008), instead of the Kingman coalescent.

Secondly, the mutation rates of many viruses, particularly RNA viruses, are among the highest observed across taxa (Lauring *et al.*, 2013; Cuevas *et al.*, 2015). Though these high rates of mutation are what enable new beneficial mutations to arise, potentially allowing for rapid resistance to host immunity or antiviral drugs, they also render high mutational loads (Sanjuán, 2010; Lauring *et al.*, 2013). Specifically, the distribution of fitness effects (DFE) has now been described across taxa-demonstrating that the input of deleterious mutations far outnumbers the input of beneficial mutations (Acevedo *et al.*, 2014; Bank *et al.*, 2014; Bernet and Elena, 2015; Jiang *et al.*, 2016). The purging of these deleterious mutants through purifying selection can affect other areas in the genome through a process known as background selection (BGS) (Charlesworth *et al.*, 1993). Accounting for these effects is important for accurate evolutionary inference in general (Ewing and Jensen, 2016), but essential for the study of viruses due to their particularly high rates of mutation and compact genomes (Renzette *et al.*, 2016).

Given these distinctive features of virus populations and the increasing use of population genetic inference in this area (*e.g.*, Renzette *et al*, 2013; Foll *et al*, 2014; Pennings *et al*, 2014; Renzette *et al*, 2016), it is crucial to account for these processes that are shaping the amount and distribution of variation across their genomes. We aim here to draw particular attention to multiple-merger coalescent events and background selection, and the repercussions of ignoring them in population genetic inference, highlighting particular applications to viruses. We conclude with general recommendations for how best to address these topics in the future.

## Skewed Offspring Distributions and the Multiple Merger Coalescent

### Inferring evolutionary history using the Wright-Fisher model: benefits and shortcomings

Many population genetic statistics and subsequent inference are based on the Kingman coalescent and the Wright-Fisher (WF) model (Wright, 1931; Kingman, 1982). With increasing computational power, the WF model has also been implemented in forward-time methods, which allows for the modeling of more complex evolutionary scenarios versus backward-time methods. This also allows for the inference of population genetic parameters, including selection coefficients and effective population sizes (*N_e_*), even from time-sampled data (i.e., data collected at successive time points) (Ewens, 1979; Williamson and Slatkin, 1999; Malaspinas *et al.*, 2012; Foll *et al.*, 2014; Foll *et al.*, 2015; Ferrer-Admetlla *et al.*, 2016; Malaspinas, 2016). These methods are robust to some violations of WF model assumptions, such as constant population size, random mating, and non-overlapping generations, and also have been extended to accommodate selection, migration and population structure (Neuhauser and Krone, 1997; Nordborg, 1997; Wilkinson-Herbots, 1998).

However, it has been suggested that violations of the assumption of a small variance in offspring number in the WF model, and in other models that result in the Kingman coalescent in the limit of large population size, lead to erroneous inference of population genetic parameters (Eldon and Wakeley, 2006). Biological factors such as sweepstake reproductive events, population bottlenecks, and recurrent positive selection may lead to skewed distributions in offspring number (Eldon and Wakeley, 2006; Li *et al.*, 2014); examples include various prokaryotes (plague), fungi (*Z. tritici, P. striiformis*, rusts, mildew, oomycetes), plants (*A. thaliana*), marine organisms (sardines, cods, salmon, oysters), crustaceans (*Daphnia*), and insects (aphids) (reviewed in Tellier and Lemaire, 2014). The resulting skewed offspring distributions can also result in elevated linkage disequilibrium (LD) despite frequent recombination, as linkage depends not only on recombination rate, but also on the degree of skewness in offspring distributions (Eldon and Wakeley, 2008; Birkner *et al.*, 2013). Such events may also skew estimates of *F*_ST_ relative to those expected under WF models, as there is a high probability of alleles being *identical by descent* in subpopulations, where the expectation of coalescent times within subpopulations is less than that between subpopulations regardless of the timescale or magnitude of gene flow (Eldon and Wakeley, 2009).

The assumption of small variance in offspring number may often be violated in virus populations as well. For example, progeny RNA virus particles from infected cells can vary up to 100 fold (Zhu *et al.*, 2009). Second, features such as diploidy, recombination, and latent stages are expected to increase the probability of multiple merger events (Davies *et al.*, 2007; Taylor and Véber, 2009; Birkner *et al.*, 2013). Third, within their life cycle, viruses experience bottleneck events during transmission and compartmentalization, followed by strong selective pressure from both the immune system and drug treatments. Finally, at the epidemic level, extinction-colonization dynamics drive population expansion (Anderson and May, 1991).

All of these aspects characterize HIV for example, a diploid virus with extraordinary rates of recombination (Schlub *et al.*, 2014). Transmitted and founder viruses undergo at least two distinct genetic bottlenecks (one of physical transmission and one of infection, respectively; Joseph and Swanstrom, 2015), followed by strong selection imposed by the immune system (Moore *et al.*, 2002). At the epidemic scale, besides multiple events of colonization (Tebit and Arts, 2011), strong heterogeneity in the virus transmission chain has also been observed (*e.g.*, Service and Blower, 1995).

### Beyond WF assumptions: the Multiple Merger Coalescent

A more general coalescent class of models, summarized as the MMC class, can account for these violations, particularly for (non-Poisson) skewed offspring distributions, by allowing more than two lineages to coalesce at a time (Table 1). These are often derived from Moran models, (Moran, 1958), generalized to allow multiple offspring per individual. In contrast to the Kingman coalescent (for which P(k>2);=0, where *k* is the number of lineages coalescing simultaneously), a probability distribution for k-merger events determines coalescence.

The parameters inferred under the MMC differ from those inferred under the Kingman coalescent in several notable respects. In a Kingman coalescent, effective size *N_e_* scales linearly with census size *N*, whereas for the MMC it does not (Huillet and Möhle, 2011). Thus genetic diversity is a non-linear function of population size. Coalescent trees under the MMC also have more pronounced star-like genealogies with longer branches (Figure 1), and their site frequency spectra (SFSs) are skewed toward an excess of low frequency and high frequency variants because of these branch lengths (Eldon and Wakeley, 2006; Blath *et al.*, 2016), generating a more negative Tajima’s *D* (Birkner *et al.*, 2013). With similar migration and population size, alleles fix at a higher rate per population in the MMC than under the Kingman coalescent, and thus higher *F*_ST_ is expected between subpopulations (Eldon and Wakeley, 2009). Further, the efficacy of selection increases, as selection acts almost deterministically between multiple merger events; in the Wright-Fisher model, genetic drift counteracts selection fairly strongly (Der et al. 2011), but in generalized models where offspring distributions are wide, beneficial mutations may be more likely to escape stochastic loss and thus continue to fixation. Furthermore, the fixation probability of a new mutant with a positive selection coefficient approaches 1 as the population size increases, in stark contrast with traditional expectations under the standard Wright-Fisher model (Der *et al.*, 2011).

Not accounting for skewed offspring distributions can lead to mis-inference. For instance, Eldon and Wakeley (2006) showed that for Pacific oysters, which have been argued to undergo sweepstake-like reproductive events (Hedgecock, 1994a), the estimated population-wide mutation rate *θ* inferred under the Kingman coalescent is two orders of magnitude larger than that obtained from the Ψ-coalescent (see below) −9 vs 0.0308, respectively-and, indeed, provides a poor fit to the data.

**Figure 1:**
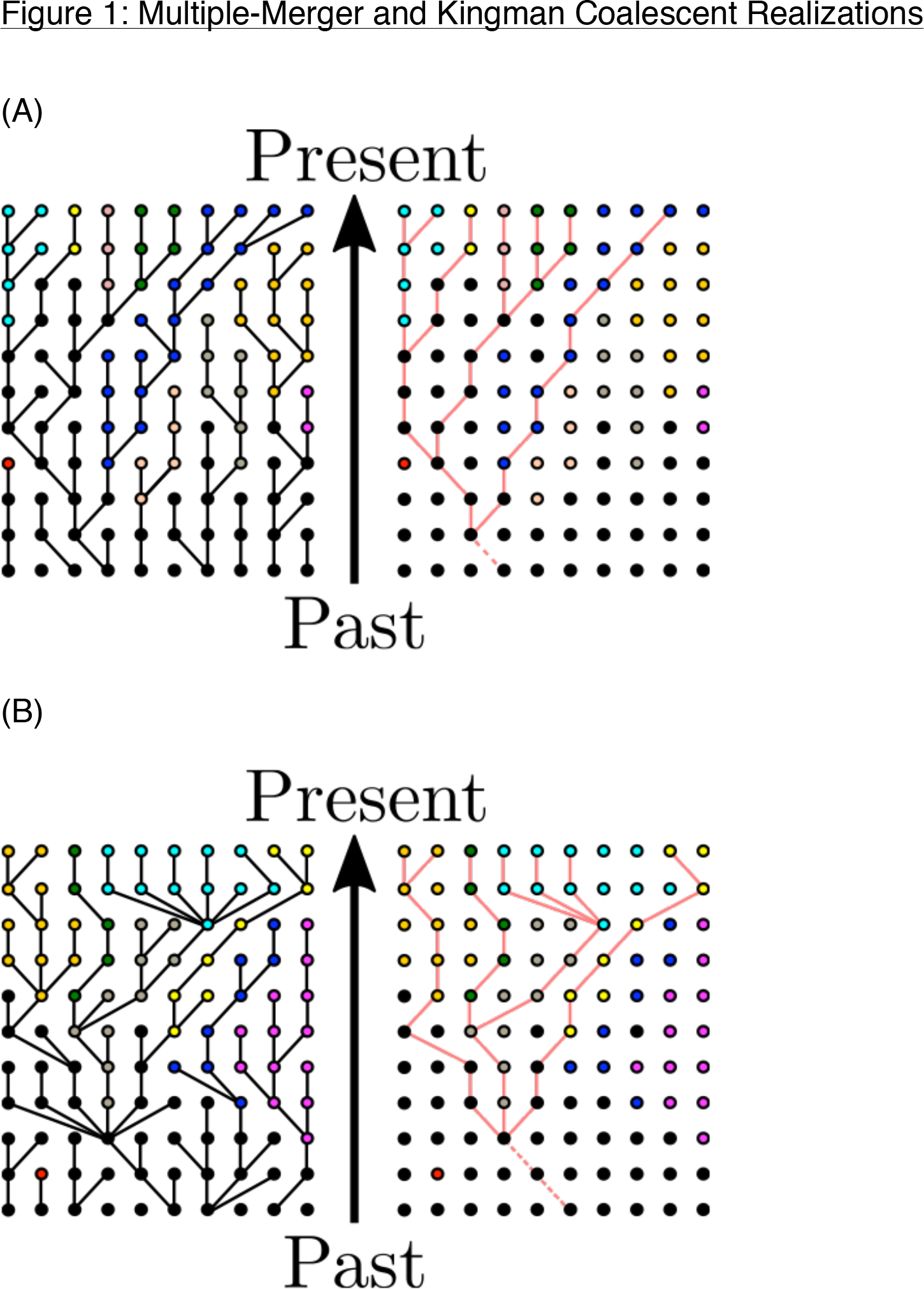
Example genealogies and samples from (A) the Kingman coalescent and (B) a multiple-merger coalescent. Panels on the left show the evolutionary process of the whole population, whereas those on the right show a possible sampling and its resulting genealogy. Colors correspond to different (neutral) derived allelic states, where black denotes the wild type.

**Table I:**
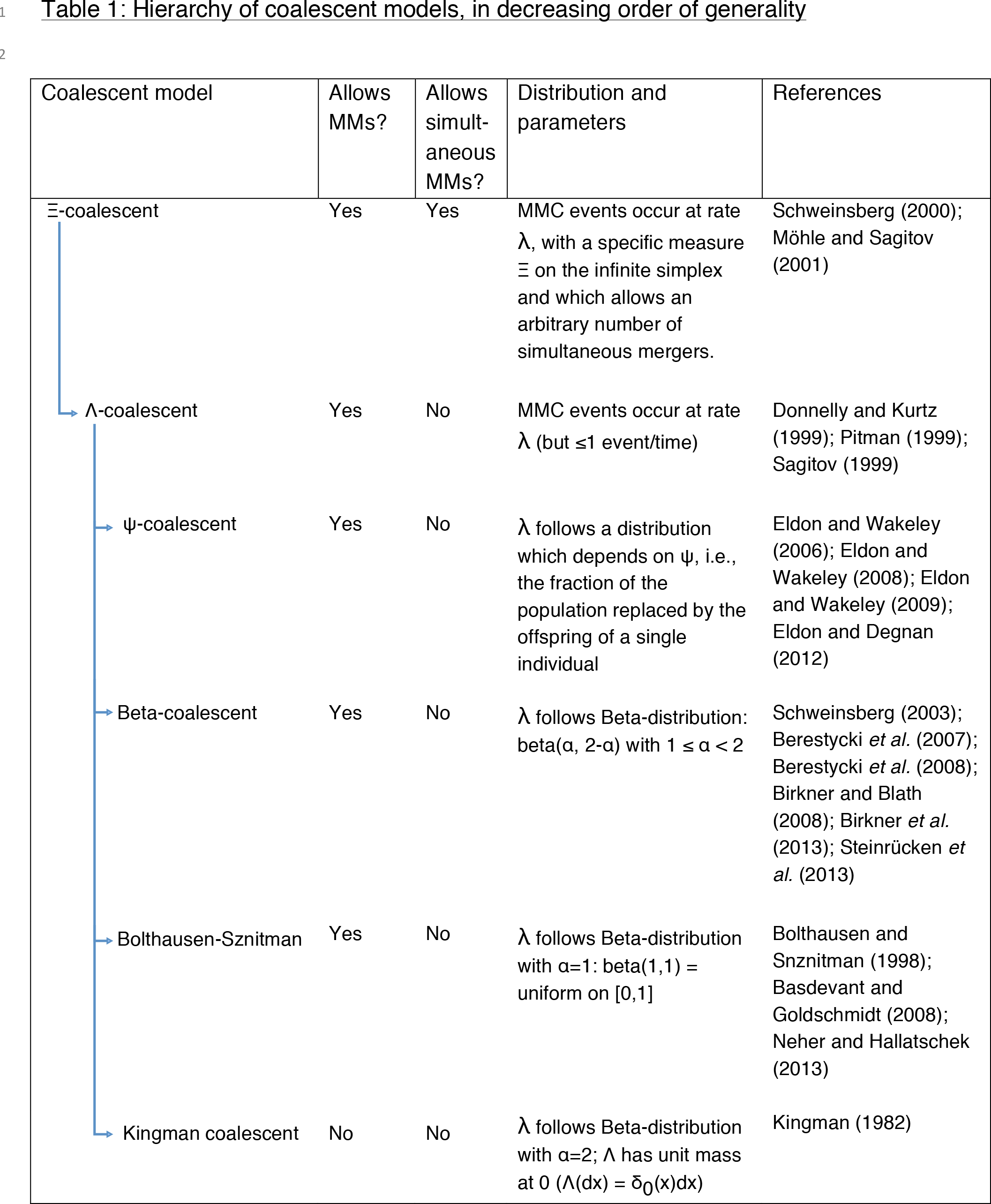
Coalescent models listed in decreasing order with respect to generality; arrows indicate coalescents that are considered subtypes of those above.

### The ψ-Coalescent

Introduced by Eldon and Wakeley (2006), the ψ-coalescent (also called the ‘Dirac-coalescent’) differentiates two possible reproductive events in the underlying forward process (Figure 2). Either a standard Moran model reproduction event occurs (with probability 1-ε), where a single individual is randomly chosen to reproduce and the (single) offspring replaces one randomly chosen non-parental individual; all other individuals, including the parent, persist. Alternatively, a ‘sweepstake’ reproductive event occurs (with probability ε) (Hedgecock, 1994b), where a single parent replaces ψ*N individuals. If these sweepstake events happen frequently enough, the rate of ψ*N-reproduction events will be much greater than that of 2-reproduction events, and the underlying coalescent process will consequently be characterized by MM events; if two or more parents were to replace ψ*N individuals, simultaneous MM events may occur in a single generation resulting in a Ξ-coalescent. However, in contrast to other MMC models (*e.g.*, Ξ-coalescent or other λ-coalescents), the parameter ψ has a clear biological interpretation as the fraction of the population that is replaced in each sweepstake reproductive event. Though the assumption of a fixed ψ (as in the normal ψ-coalescent) seems biologically unrealistic, it can be avoided by treating ψ as a Poisson parameter. Finally, despite its appealing connection to biologically relevant measures, the appropriateness of making inferences based on the ψ-coalescent still depends on the biology of the specific virus being studied. Thus, model choice is still essential, and the best-fit coalescent should be assessed on a case-by-case basis.

**Figure 2:**
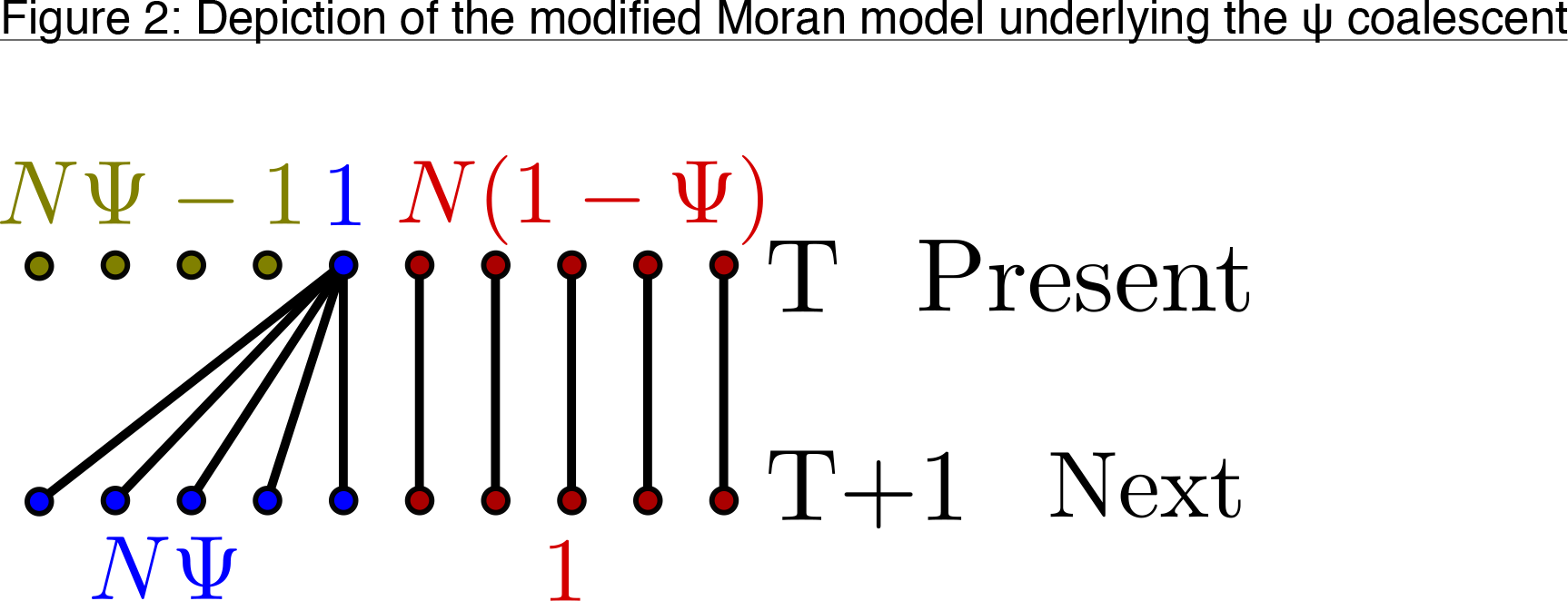
Lineages between the present and the next generation, where *N* is the population size, ε is the probability of a sweepstake event, and ψ is the fraction of the population that is replaced in each such event. Labels in the top row give the number of parental individuals reproducing in a given manner (represented by color), whereas labels in the bottom row give the number of corresponding offspring per parent.

### Application to Viruses

There are several reasons why a modified Moran model may better capture virus evolution than models converging to the Kingman coalescent, although it does not account for fitness differences between individuals. First, virus evolution is driven by strong bottlenecks during host transmission and intrahost selection processes, which likely result in skewed offspring distributions (Figure 3) (Gutiérrez *et al.*, 2012; Tellier and Lemaire, 2014). Further, viruses display the MMC-typical low *N_e_/N* ratio (Pennings *et al.*, 2014; Tellier and Lemaire, 2014), can adapt rapidly (Neher and Hallatschek, 2013), and may have sweepstake-like reproductive events in which a single virion can propagate a large fraction of the entire population (Grenfell *et al.*, 2004; Pybus and Rambaut, 2009). For example, the influenza virus hemagglutinin (HA) segment appears to be under strong directional selection imposed by host immunity (and sometimes drug treatment), resulting in a ladder-like genealogy, (as depicted in Figure 3A), suggesting that only a few viruses seed the entire next generation (Grenfell *et al*, 2004). That being said, some challenges remain, such as rigorously defining the term ‘generation’ for virus populations, and subsequently confirming that the per generation mutation rate is on the order of the coalescent timescale c_N_, which is a prerequisite for the use of any coalescent approach. Finally, viruses with little or no recombination may be prone to clonal interference, which should be explicitly accounted for in population models and resulting coalescents *(e.g.*, Strelkowa and Lässig, 2012).

Those processes that make viruses ideal candidates for MMCs can differ by scale (see Figure 3); for example, following transmission events, there are severe founder events and potentially high recombination within the host (*e.g.*, HIV, HCMV). Subsequent compartmentalization may introduce intra-host population structure through bottlenecks, colonization events, and extinction events (Renzette *et al.*, 2013). To date, it remains unclear how often MMCs fit the patterns of variation observed in intra-host relative to inter-host virus populations-but such comparisons are increasingly feasible. Finally, periods of latency-temporary virus inactivation with cessation of reproduction-should be incorporated in such modeling, potentially as recurring mass extinction events (Taylor and Véber, 2009). Thus, multiple MMC models are a necessary but not final step towards addressing the various patterns observed at different scales of virus evolution (Table 1).

The large data sets often generated from viruses may also prove impractical for the likelihood-based methods commonly employed for MMCs. This limitation has partially been overcome by Eldon *et al.* (2015), who proposed an approximate likelihood method along with an Approximate Bayesian Computation (ABC) approach based on the SFS to distinguish between the MMC and exponential population growth. Although both effects are expected to result in very similar SFSs, characterized by an excess of singletons as compared to the Kingman coalescent, the bulk and tail of the SFS (i.e., the higher-order frequency classes) typically differ, which can be assessed by approximate likelihood-ratio tests and Approximate Bayes Factors (Eldon *et al.*, 2015).

**Figure 3:**
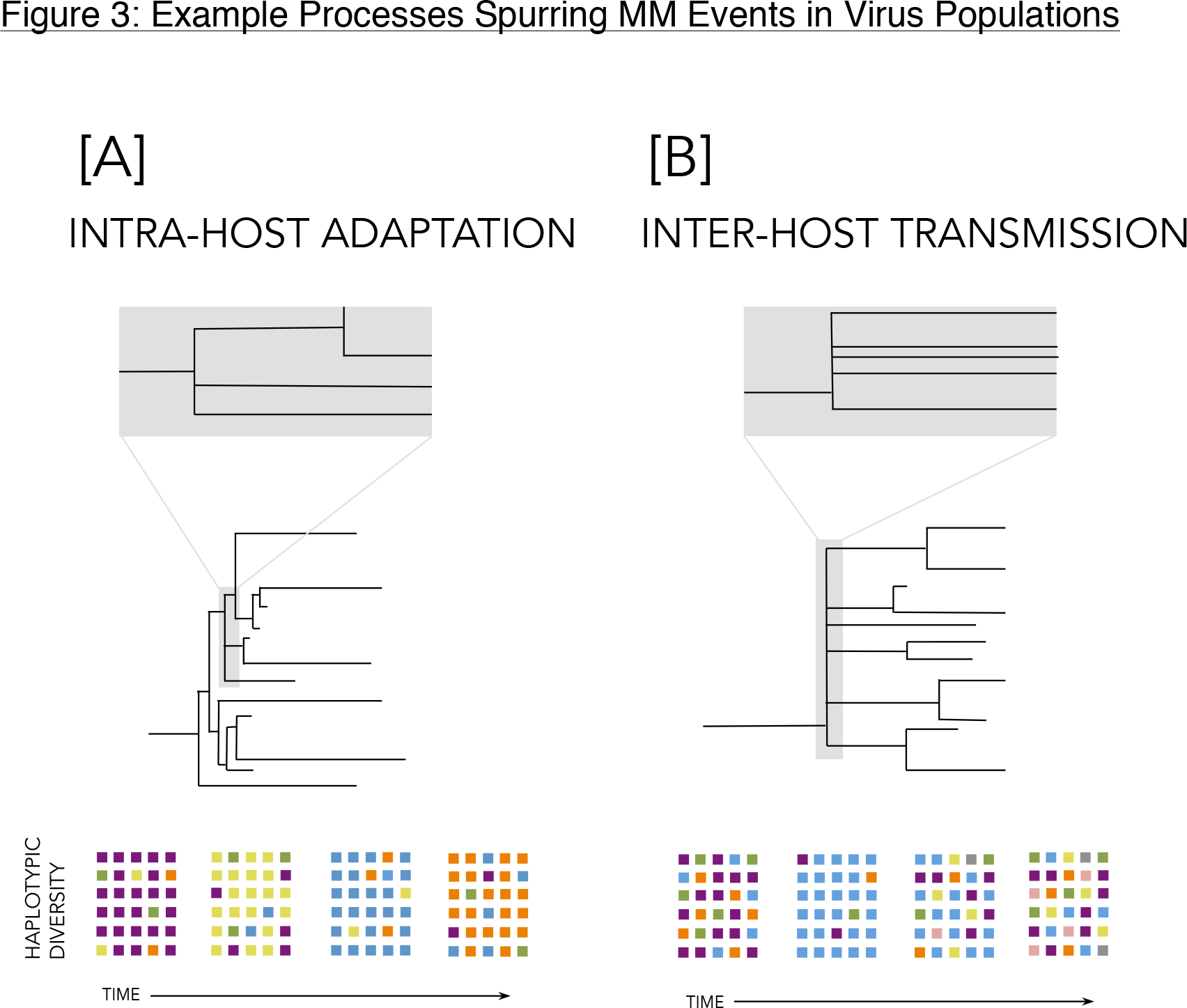
Examples include (A) intra-host adaptation (a selective process) and (B) inter-host transmission (a demographic process). The tree in (A) characterizes, for example, NA or HA evolution in the influenza A virus, driven by positive selection; selection by host immunity is ongoing, while that from drug treatment may be intermittent. The tree in (B) represents inter-host transmission and its associated bottleneck; for viruses that compartmentalize (such as HCMV and HIV), similar patterns follow transmission to new compartments. The colored squares below the trees roughly indicate the diversity of the population through time. Intra-host adaptation may temporally decrease diversity owing to genetic hitchhiking, though single snapshots may not reflect varying temporal levels of diversity. During inter-host transmission, diversity decreases owing to the associated bottleneck but then may quickly recover in the new host.

### BOX 1: Future Challenges in MMC Models

#### In order to make MMC models biologically relevant for viruses, a number of important tasks remain

1. Describe summary statistics that capture demographic features and processes when offspring distributions are highly skewed; such patterns will be required for large-scale inference in a computationally efficient (*e.g.*, Approximate Bayesian) framework.
2. Better understand the behavior of commonly used summary statistics under such models, as done for *F*_ST_ by Eldon and Wakeley (2009), for commonly used divergence, SFS, and LD-based statistics.
3. Determine which MMCs are best suited for different scales of virus evolution (i.e., intra-host, inter-host, global); develop novel models if necessary.
4. Investigate the effect of violations of MMC assumptions (*e.g.*, overlapping generations, number of multiple merger events) on inference.

[END BOX 1]

## Purifying Selection and Linkage in Virus Populations

### Modeling Background Selection

The joint modeling of the effects of genetic drift and positive selection, including in experimental evolution studies of virus populations, has improved our ability to distinguish adaptive from neutral mutations by minimizing the chance that the rapid fixation of a neutral allele is incorrectly interpreted as strong positive selection (Li *et al.*, 2012; Foll *et al.*, 2014). However, there is another process that must be incorporated if we are to fully understand mutation trajectories in virus populations: background selection (BGS).

BGS was originally proposed to explain patterns of reduced diversity in regions of low recombination-patterns that were previously suggested to be the signature of genetic hitchhiking (HH) around strongly beneficial mutations (see Begun and Aquadro, 1992 and Charlesworth *et al.*, 1993). It was argued that only neutral mutations present on the “least-loaded” chromosomes-that is, those with the fewest deleterious mutations – have appreciable probabilities of reaching high frequencies or fixation. Kimura and Maruyama (1966) showed that the proportion of chromosomes belonging to the least-loaded class is

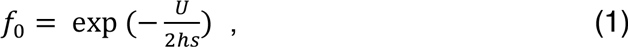

where *U* is the rate of mutation to a deleterious state, s is the selection coefficient against homozygous mutations, and *h* is the dominance coefficient. For simplicity of modeling, *h* is usually set to 1 for viruses that carry a single copy of their genome in each virion, although polyploid effects could arise in the case of multiple virions infecting the same cell.

The least-loaded class, and thus genetic diversity in the presence of BGS, is dependent on the balance between the influx of deleterious mutations (occurring at rate *U*) and their removal by natural selection (according to the product hs). Assuming that offspring exclusively originate from the least-loaded class of individuals, Charlesworth *et al.* (1993) expressed the expected neutral diversity due to background selection as

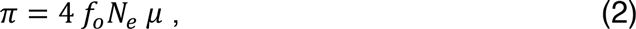

where *N_e_* is the effective population size and *μ* is the mutation rate. As BGS reduces the number of reproducing individuals, genetic drift increases, thus reducing genetic diversity and increasing stochasticity in allele trajectories. Further, since only the genetic diversity segregating in the least-loaded class can be observed, population size inferred from measures of genetic diversity may be underestimated if BGS is not properly taken into account (Ewing and Jensen, 2016).

In the BGS model described above, strongly deleterious mutations are maintained in mutation-selection balance such that no skew in the SFS is expected, as rare variants are rapidly purged. Thus, a simple re-scaling of *N_e_* is often used as a proxy for the effects of BGS (*e.g.*, Hudson and Kaplan, 1995; Zeng and Charlesworth, 2011; Prüfer *et al*, 2012; Zeng, 2013). However, recent work has demonstrated that, while this re-scaling is appropriate for strongly deleterious mutations, it is largely inappropriate for weakly deleterious mutations that may segregate in the population. Figure 4 shows the skew in estimates of population size and migration rates obtained using an ABC approach when BGS is prevalent for two populations A and B that have split at time *τ=2N_e_* generations (reproduced from Ewing and Jensen,2016). Further, experimental work on the shape of the distribution of fitness effects (DFE) in many organisms indicates that weakly deleterious mutations represent an important class (e.g., Eyre-Walker and Keightley, 2007; Bank *et al*, 2014). These mutations may act to skew the SFS towards rare alleles as they decrease the expected frequency of linked neutral mutations relative to neutral expectations. As subsequent demographic inference is based on the shape of this SFS, this effect should be properly accounted for by directly simulating weakly deleterious mutations rather than implementing a simple rescaling, as is common practice. Though important analytical progress has been made in this area (*e.g.*, McVean and Charlesworth, 2000), simulations remain the best option for the non-equilibrium demographic models and alternative coalescents recommended here for inference in virus populations.

**Figure 4:**
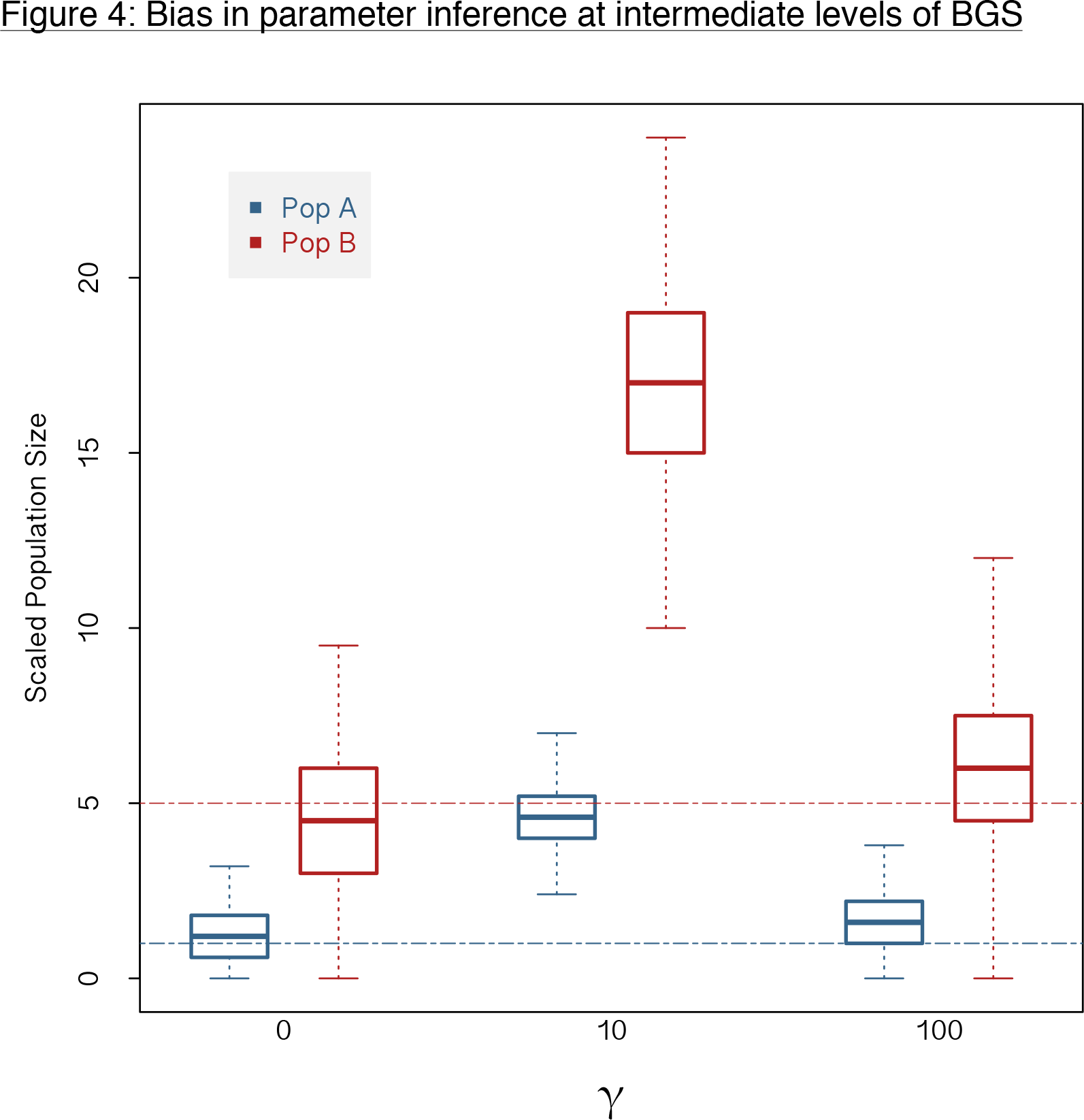
Bias in parameter inference for different levels of BGS, redrawn from Ewing & Jensen (2016). Posterior densities from ABC inference for population size are shown. The strength of purifying selection is given as *γ*, where *γ*;=;*2NeS.* Population A has a true scaled size of 1 (blue line), and population B a true scaled size of 5 (red line). Both population sizes are scaled relative to the size of the ancestral population. As shown, the greatest mis-inference occurs in the presence of weakly deleterious mutations and subsequent strong BGS effects.

### The Effects of Background Selection on Inference in Virus Populations

Efforts to estimate the impact of BGS in non-viral organisms have been well reported. One of the most notable examples is that of Comeron (2014), who estimated levels of BGS in *Drosophila melanogaster* based on the results of Hudson and Kaplan (1995) and Nordborg *et al.* (1996) using a high-definition recombination map, with results indicating strong effects across the genome. For viruses, similar efforts are in their infancy, with the first attempt at such estimation in a virus reported recently by Renzette *et al.* (2016), utilizing the theoretical predictions of Innan and Stephan (2003). Interestingly, the full spectrum of recombination frequencies is available in viruses – from nonrecombining (*e.g.*, most negative-sense RNA viruses), to re-assorting (*e.g.*, Influenza virus), to rarely recombining (*e.g.*, Hepatitis C and West Nile viruses), to frequently recombining (e.g., HIV), offering a highly promising framework for comparative analyses investigating the pervasiveness of BGS effects (Chare *et al.*, 2003; Simon-Loriere and Holmes, 2011). Further, given the high mutation rates and compact genomes of many viruses, evolutionary theory suggests effects at least equal to those seen in *Drosophila*.

In order to accomplish such inference, improved recombination maps for virus genomes will be important. With such maps in hand, and given the amenability of viruses to experimental perturbation, it may indeed be feasible to understand and account for BGS in models of virus evolution.

### BOX 2: Future challenges in identifying the effects of BGS

#### As BGS almost certainly impacts inference in virus populations, accounting for its effects is critical. Future challenges include

1. Account for BGS effects on the SFS by directly simulating weakly deleterious mutations, rather than by rescaling *N_e_*.
2. Improve recombination maps for virus genomes.
3. Develop models combining the effects of non-equilibrium demography, positive selection, and BGS, ideally to allow for the joint estimation of all associated parameters.
4. Extend methods applied to other taxa to virus populations; for example, establishing a baseline of variation for use as a null expectation to estimate BGS levels across the genome, as done for *Drosophila*.

[END BOX 2]

## Future Directions

Given that skewed offspring distributions and pervasive linked selection are likely important factors influencing the inference of virus population parameters, it is important to note that multiple backward and forward simulation programs have recently been developed which make the modeling of these processes feasible (Hernandez, 2008; Messer, 2013; Thornton,2014; Eldon *et al.*, 2015; Zhu *et al.*, 2015). This will allow researchers to directly simulate from parameter ranges that may be relevant for their population of interest, developing a better intuition for the importance of these processes in shaping the observed genomic diversity. More concretely, the ability to now simulate in a computationally efficient framework opens the possibility of directly implementing ABC inference approaches under these models. Thus, by drawing mutations from a biologically realistic distribution of fitness effects and allowing offspring distributions to appropriately vary, it is now possible to re-implement common demographic estimation or genome scan approaches; these modified approaches would be based on more appropriate null expectations of the shape of the SFS, the extent of linkage disequilibrium, and the degree of population divergence.

## Acknowledgments

We would like to thank Bjarki Eldon for helpful suggestions during the early stages of this manuscript. This work was funded by a European Research Council (ERC) Starting Grant to JDJ, as well as Swiss National Science Foundation (FNS) grants to JDJ (31003A_159835) and SV (PMPDP3_158381).

## Data Archiving

As a review article, no new data was processed, analyzed, or used directly.

